# Cysteine post-translational modifications regulate protein interactions of caveolin-3

**DOI:** 10.1101/2022.09.15.508083

**Authors:** Fiona Ashford, Elaine Brown, Sarah Calagan, Izzy Jayasinghe, Colin Henderson, William Fuller, Krzysztof Wypijewski

## Abstract

Caveolae are small flask-shaped invaginations of the surface membrane which are proposed to recruit and co-localise signalling molecules. The distinctive caveolar shape is achieved by the oligomeric structural protein caveolin, of which three isoforms exist. Aside from the finding that caveolin-3 is specifically expressed in muscle, functional differences between the caveolin isoforms have not been rigorously investigated. Caveolin-3 is relatively cysteine-rich compared to caveolins 1 and 2, so we investigated its cysteine post-translational modifications. We find that caveolin-3 is palmitoylated at 6 cysteines and becomes glutathiolated following redox stress. We map the caveolin-3 palmitoylation sites to a cluster of cysteines in its C terminal membrane domain, and the glutathiolation site to an N terminal cysteine close to the region of caveolin-3 proposed to engage in protein interactions. Glutathiolation abolishes caveolin-3 interaction with heterotrimeric G protein alpha subunits. Our results indicate that a caveolin-3 oligomer contains up to 66 palmitates, compared to up to 33 for caveolin-1. The additional palmitoylation sites in caveolin-3 therefore provide a mechanistic basis by which caveolae in smooth and striated muscle can possess unique phospholipid and protein cargoes. These unique adaptations of the muscle-specific caveolin isoform have important implications for caveolar assembly and signalling.

## Introduction

Caveolae are small, flask shaped invaginations of the cell surface membrane that recruit and co-localise receptors, G proteins, their effector molecules and downstream targets to ensure fidelity of signal transduction in numerous tissues (1-3). Classically, the predominant structural component of caveolae was thought to be the scaffolding protein caveolin, which recruits both lipids and proteins to assemble the caveolar architecture (4,5). Three caveolin isoforms exist: caveolins 1 and 2 are ubiquitous, and caveolin-3 expression is confined only to smooth and striated muscle (6). The cryo-EM structure of recombinant caveolin-1 reveals it forms a flat, disk-shaped oligomer composed of 11 monomers (7). In recent years a second family of obligatory caveolar structural proteins, the cavins, has been identified. Cavins stabilise the caveolin oligomer and like caveolins are obligatory for the formation of caveolae (8-10). Cavin-1 is ubiquitous and indispensable for caveolar formation (11). Cavins 2-4 show more tissue-specific distributions.

The dynamic association of proteins with caveolae offers an opportunity for regulation of signal transduction, but remarkably few proteins have been found to be recruited into or expelled from caveolae, and the caveolar proteome remains extremely stable during signal transduction (12). Post-translational modifications of caveolins and cavins represent one means by which their protein interactions could be regulated. For example, phosphorylation of caveolin-1 Tyr-14 by Src kinase modifies its interactions with a matrix metalloprotease (13). SUMOylation of caveolin-3 modifies β2-but not β1-adrenoceptor dependent signalling (14). Although there are numerous phosphorylation sites annotated in the cavin isoforms (8), few other post-translational modifications have been found to regulate the behaviour of cavins or caveolins. Caveolin-1 is palmitoylated at 3 cysteine residues close to its integral membrane domain (Fig 1A), but caveolin-1 palmitoylation is irreversible, and not required for caveolin-1 membrane-targeting (15,16). Analogous cysteines exist in caveolins 2 and 3 (Fig 1B), but their palmitoylation has not been rigorously investigated.

**Figure 1:**
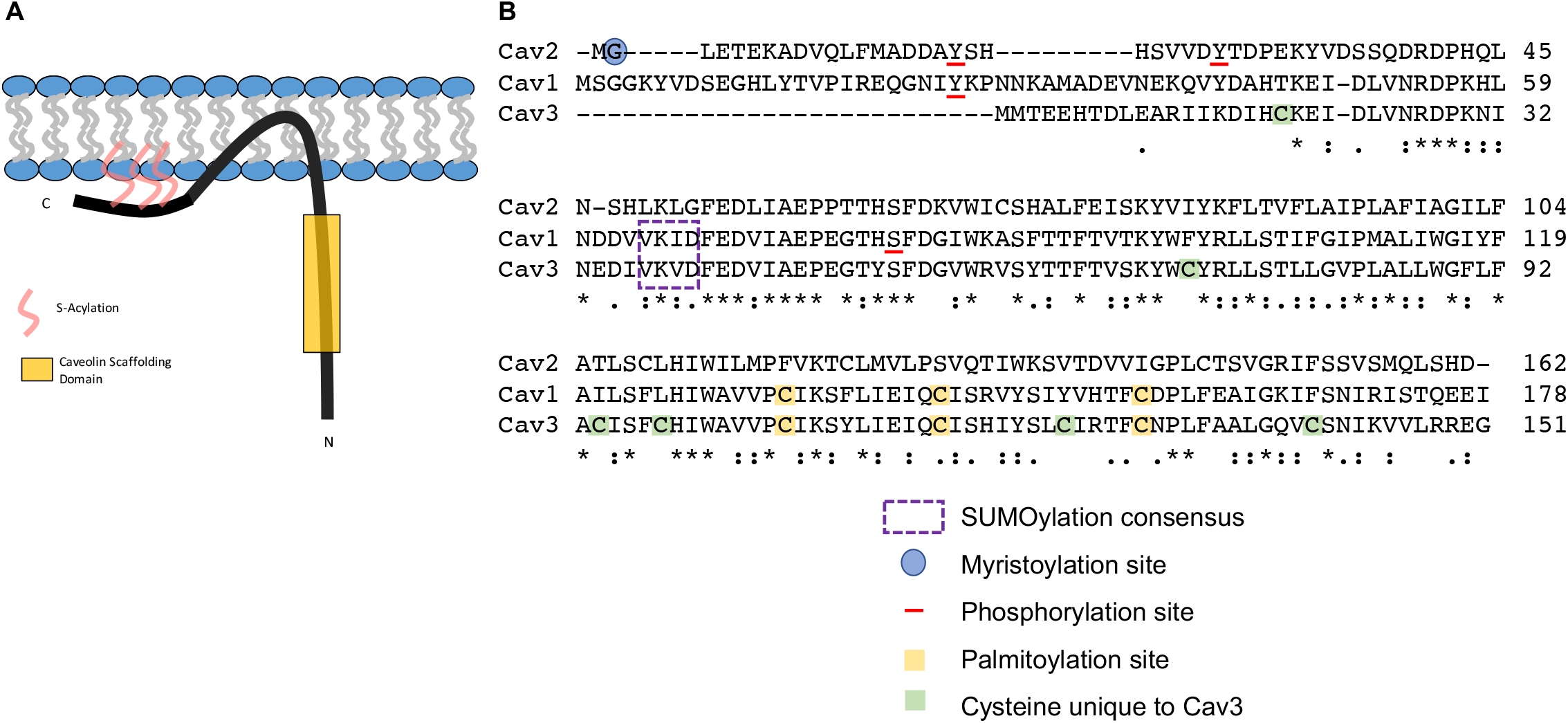
Caveolin schematic and pot-translational modifications. A: caveolin schematic, showing the positions of the proposed palmitoylation (acylation) sites and the caveolin scaffolding domain. B: Alignment of the three rat caveolin isoforms (Uniprot P41350, Q2IBC5 and P51638). The positions of the caveolin-1 palmitoylation sites, and the analogous cysteines in caveolin 3 are highlighted in yellow. The positions of the caveolin-2 myristoylation site, and experimentally-determined phosphorylation sites are also indicated. Cysteines unique to caveolin-3 are highlighted in green.

In the present investigation we characterised cysteine post-translational modifications of caveolin-3, the muscle-specific caveolin isoform. Using a novel PEGylation assay we find up to six caveolin-3 cysteines are palmitoylated in cardiac muscle and map these palmitoylation sites in transfected cells. We also find that caveolin-3 is reversibly glutathiolated, and that caveolin-3 glutathiolation modifies its interaction with heterotrimeric G proteins during redox stress. Our findings highlight important functional differences between caveolin isoforms and assign unique regulatory functions to caveolae in smooth and striated muscle.

## Results

### Palmitoylation site mapping in caveolin-3

Caveolin-1 (which has only 3 cysteines in its sequence, Fig 1B) is triply palmitoylated (15), but the palmitoylation stoichiometry of caveolin-3 has not been investigated. By exchanging palmitates for a PEG molecule, we investigated the stoichiometry of caveolin-3 palmitoylation in multiple cell types. Our acyl-PEG exchange assay generates a ∼5kDa band shift for each palmitoylation site occupied. In both the myoblast cell line H9c2 and adult rat ventricular myocytes we identified multiple palmitoylation sites in caveolin-3: at least 5 sites in H9c2 cells (Fig 2A), and 6 sites in rat ventricular myocytes (Fig 2B).

**Figure 2:**
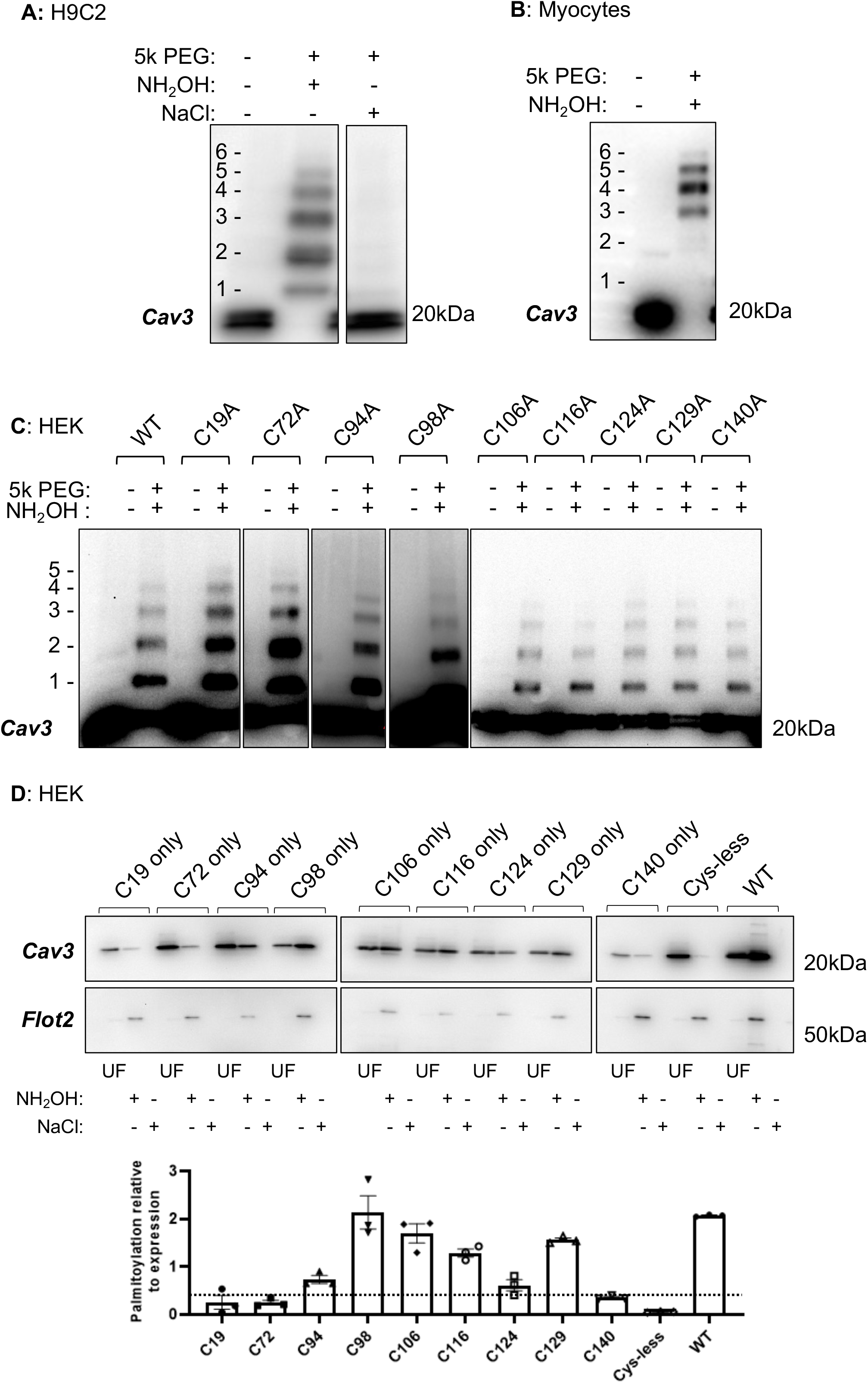
Caveolin-3 palmitoylation in cells and tissue. A: Acyl PEG exchange identifies up to 6 palmitoylation sites in caveolin-3 in H9c2 cells, with the predominant species doubly or triply palmitoylated. B: Acyl PEG exchange identifies up to 6 palmitoylation sites in caveolin-3 in rat ventricular myocytes, with the predominant species quadruply palmitoylated. C: Single cys to ala mutagenesis fails to definitively identify caveolin-3 palmitoylation sites in transfected HEK cells, where the predominant form of caveolin-3 is singly or doubly palmitoylated. D: Acyl-resin assisted capture to identify palmitoylation sites in caveolin-3. Caveolin-3 mutants with only a single cysteine intact were expressed in HEK cells and purified using acyl resin-assisted capture. Flot2: flotillin-2, UF: unfractionated cell lysate, Resin-assisted capture in the presence of hydroxylamine (NH_2_OH) or sodium chloride (NaCl, negative control) is shown. The bar chart shows the amount purified using acyl-resin assisted capture relative to expression for each caveolin-3 mutant. The dotted line on the bar chart illustrates an arbitrary threshold cutoff to identify the six most highly palmitoylated cysteines in caveolin-3.

To identify the palmitoylated cysteines in caveolin-3 we first generated single cysteine to alanine mutations and expressed them in HEK cells, which lack endogenous caveolae but possess a full repertoire of zDHHC-PATs (17). The acyl PEG exchange assays reveal that wild type caveolin-3 was significantly less palmitoylated in HEK cells than ventricular muscle (predominant palmitoylated species +1 to +2 palmitates, compared to +4 palmitates in ventricular muscle, Fig 2C). This precluded definitive identification of the caveolin-3 palmitoylation sites using this approach. We therefore generated a cys-less caveolin-3, reintroduced individual cysteines to the cys-less protein, and expressed these proteins in HEK cells. Resin-assisted capture purifies palmitoylated proteins regardless of how many palmitoylation sites are occupied (18), so is a suitable approach to assess the palmitoylation status of proteins with only one cysteine intact. All caveolin-3 mutants containing a single cysteine were palmitoylated to some extent (Fig 2D). The principal palmitoylation sites detected in caveolin-3 in these experiments were cysteines 98, 106, 116 and 129. We also detected significant palmitoylation of cysteines 94 and 124, but very little palmitoylation of cysteines 19, 72 and 140 (Fig 2D). Since the quadruple palmitoylated species is the most common detected by acyl-PEG exchange in ventricular muscle (Fig 2B), we conclude this represents caveolin-3 palmitoylated at cysteine 98, 106, 116 and 129. We propose that the additional two palmitoylation sites occupied in ventricular muscle are cysteines 94 and 124, but we do not rule out residual palmitoylation of other cysteines in caveolin-3.

### Position of the palmitoylated cysteines in the caveolin oligomer

The caveolin-1 oligomer forms a flat disk 140 Å in diameter, in which the protomers’ C termini are at the centre of the disk, and its N termini on the edge (7). We mapped the positions of the caveolin-3 cysteines in the caveolin-1 cryo-EM structure PDB-7SC0. C140 (analogous to F167 in caveolin-1), which is not palmitoylated, lies close to the centre of the disk facing the membrane (cyan in Fig 3). Cysteines 106 116 and 129 (yellow in Fig 3), which are conserved in caveolin-1, are located in two concentric circles in the oligomer, diameters approx. 50 Å and 90 Å: the inner circle is formed by C129 (C156 in caveolin-1), and the outer circle is formed from C116 (C143 in caveolin-1) from one protomer in close proximity to C106 (C133 in caveolin-1) from the neighbouring protomer. The additional palmitoylation sites in caveolin-3 (orange in Fig 3) form two additional concentric circles, diameters approx. 65 Å and 110 Å. The smaller of these, midway between the caveolin-1 palmitoylation sites, is formed by C124 (Y151 in caveolin-1). The larger is formed from C94 (I121 in caveolin-1) and C98 (L125 in caveolin-1) from the same protomer. C72 (F99 in caveolin-1, cyan in Fig 3) which is not palmitoylated lies on the outer edge of the disk, and C19 is not resolved. None of the cysteines or their analogous residues are visible on the cytosolic face of the caveolin-1 oligomer: they all face the membrane.

**Figure 3:**
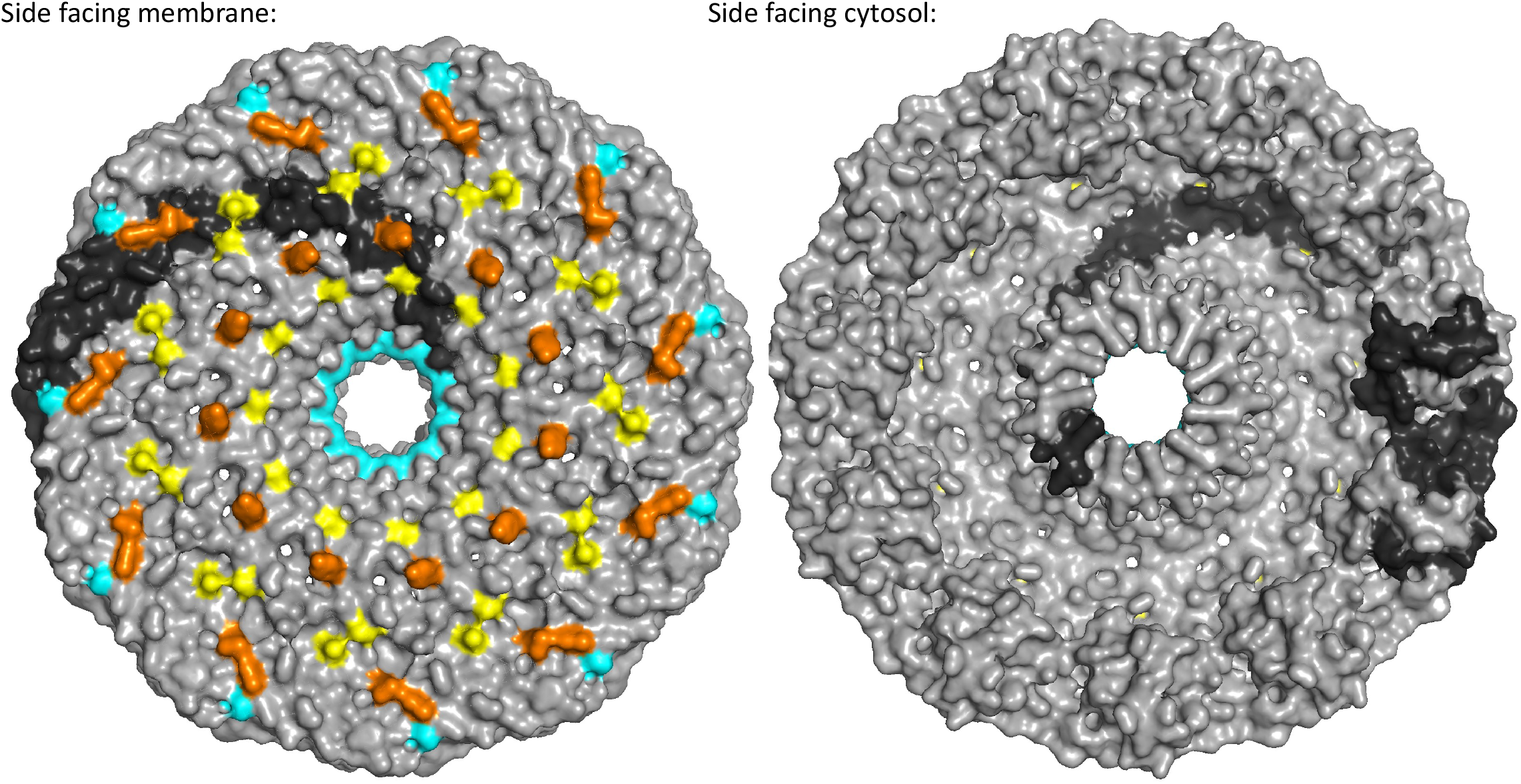
Caveolin palmitoylation sites. Positions of the caveolin-3 palmitoylation sites in the caveolin-1 oligomer PDB-7SC0. The oligomer is shown in surface representation with a single protomer in black. Cyan: caveolin-3 cysteines that are not palmitoylated. Yellow: palmitoylated cysteines shared between caveolin-1 and caveolin-3. Orange: palmitoylated cysteines unique to caveolin-3.

### Palmitoylation and protein interactions of caveolins

Caveolin forms homo-oligomers composed of 11 protomers. Our palmitoylation site mapping experiments imply that caveolin-3 homo-oligomers would incorporate up to 66 palmitate molecules. We investigated whether protein interactions of caveolin were influenced by its palmitoylation status, and whether they were different for the more heavily palmitoylated caveolin-3. We generated a caveolin-3 mutant in which only the cysteines analogous to caveolin-1 cysteines were intact (C106, C116, C129), which we denoted ‘caveolin-1-like’ (C1L). We compared the protein interactions of cysless caveolin-3, C1L and wild type caveolin-3 co-transfected with GNAI2 into HEK cells. Co-immunoprecipitation of GNAI2 with C1L or WT caveolin-3 was essentially identical (Fig 4). However, cys-less caveolin-3 co-purified significantly less with GNAI2 than either WT or C1L. Hence although non-palmitoylated caveolins are capable of oligomerising and forming the characteristic caveolar bulb (19), they do not fully support physical interactions with caveolar resident proteins. However, the additional palmitoylation sites in caveolin-3 do not influence its interaction with GNAI2 compared to caveolin-1.

**Figure 4:**
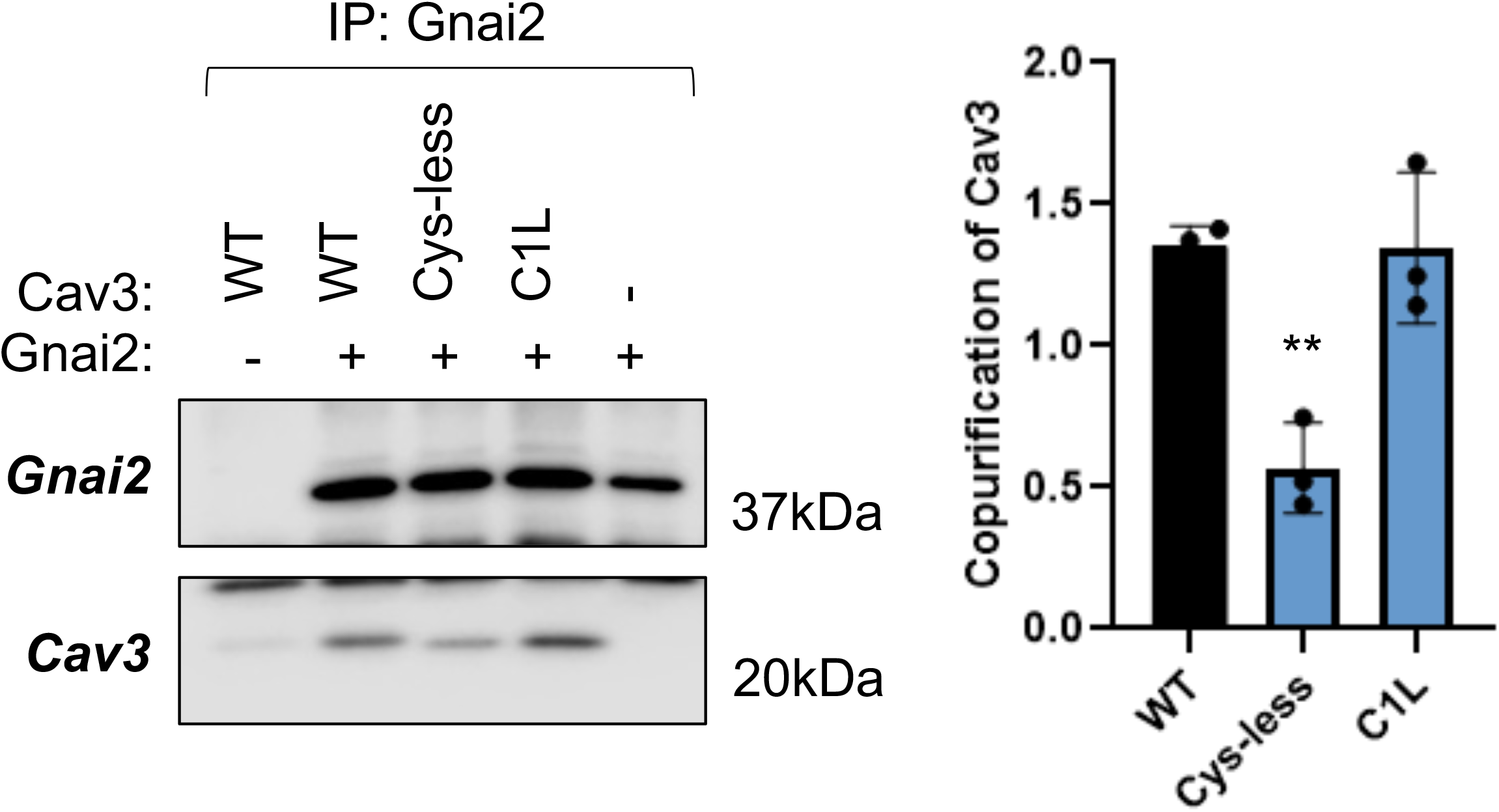
Influence of palmitoylation on caveolin protein-protein interactions. Caveolin-3 and Gnai2 were co-expressed in HEK cells, Gnai2 was immunoprecipitated and co-purification of caveolin-3 assessed. Significantly less cys-less caveolin-3 copurifies with Gnai2 than wild type (WT) or caveolin-3 with only the caveolin-1 palmitoylation sites intact (C1L). The graph shows co-purification of caveolin-3 normalised to the quantity of Gnai2 immunoprecipitated (n=3; **: P<0.01 compared to WT, Dunnett’s multiple comparisons test).

### Glutathiolation of caveolin-3

We loaded rat ventricular myocytes with cell permeable biotinylated glutathione ethyl-ester (bioGEE), purified glutathiolated proteins using streptavidin beads and identified them using mass spectrometry (Fig 5A). We considered proteins to be bone fide glutathiolation candidates if they were detected in three out of four independent experiments (Fig 5B), which generated a list of 150 glutathiolated proteins in rat ventricular myocytes (Supplemetary Table 1). We validated candidate glutathiolated proteins by immunoblotting (Fig 5C) and confirmed that streptavidin capture of selected candidates (Gαs and caveolin-3) required myocytes to be loaded with biotinylated glutathione ethyl ester (Fig 5D). Subsequent experiments focussed on glutathiolation of caveolin-3 in ventricular muscle.

**Figure 5:**
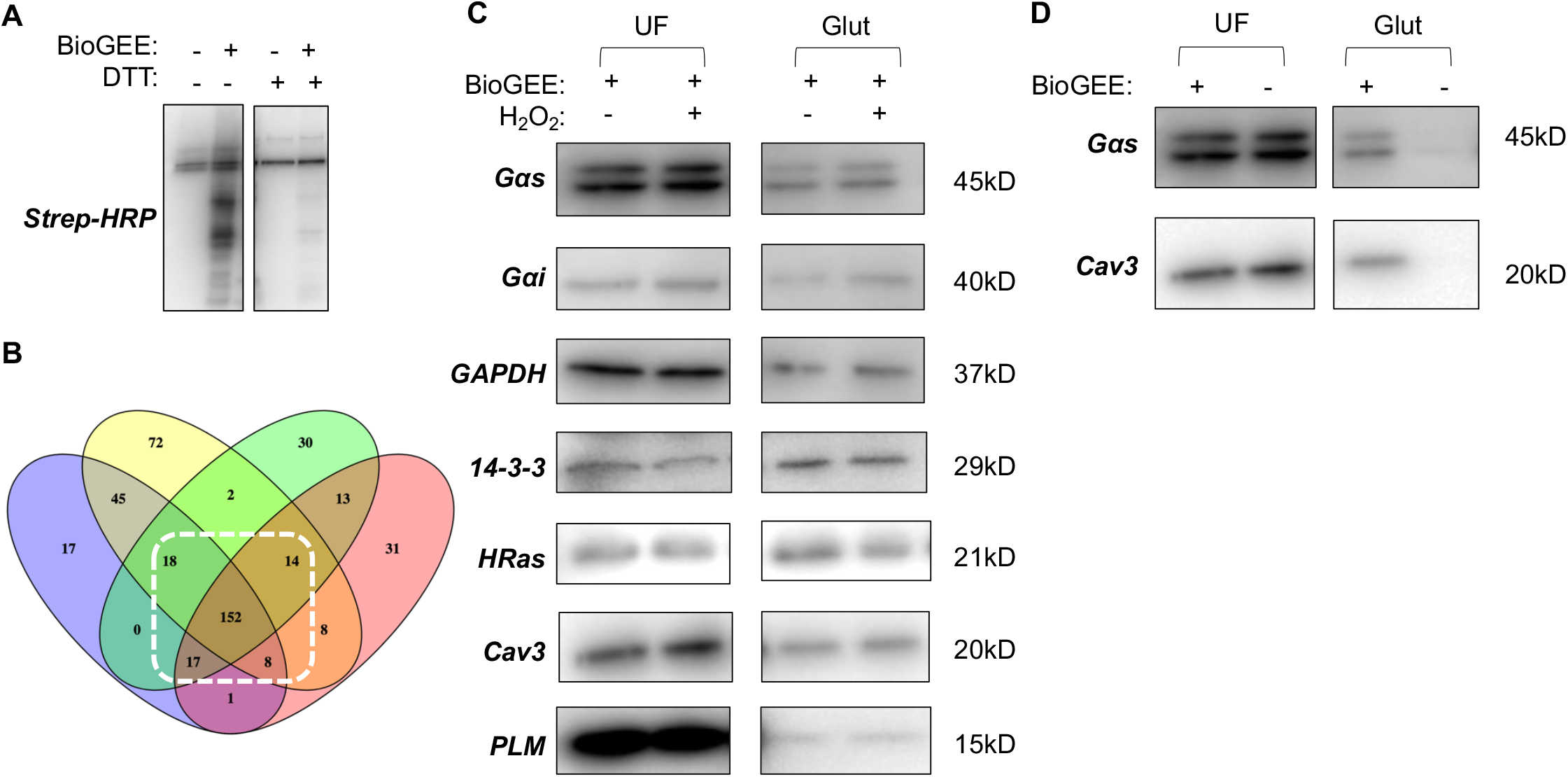
Identification of glutathiolated proteins in rat ventricular myocytes. A: biotin incorporation into cells loaded with biotinylated glutathione ethyl ester (bioGEE) is sensitive to the reducing agent DTT. B: Proteomic analysis of glutathiolated proteins purified from rat ventricular myocytes identifies 209 high confidence glutathiolated proteins (detected in at least 3/4 experiments). C: Validation of selected glutathiolated proteins identified by mass spectrometry (UF: unfractionated cell lysate, Glut: purified glutathiolated proteins). D: Purification of Gαs and caveolin-3 using the bioGEE protocol is dependent on preloading cells with bioGEE.

We detected baseline glutathiolation of caveolin-3 in ventricular myocytes, which was elevated following treatment with the thiol stressor diamide but not by application of the exogenous oxidant hydrogen peroxide, the NADPH oxidase agonist angiotensin II, or oxidised gluthathione disulphide (Fig 6A). In separate experiments we sought to identify reversibly oxidised cysteines in caveolin-3 by replacing hydroxylamine with DTT in our PEGylation assay. Treatment of ventricular myocytes with diamide substantially increased PEGylation of caveolin-3 at a single cysteine in this assay (Fig 6B). Since diamide is generally agreed to generate a thiol stress without generating free radicals (20,21), we interpret this band shift to indicate that caveolin-3 is predominantly glutathiolated at a single cysteine in ventricular myocytes. To identify this glutathiolation site we expressed single cysteine mutants of caveolin-3 in HEK cells and treated the cells with diamide. In our panel of single cys to ala mutants of caveolin-3, no single C to A mutation significantly reduced caveolin-3 glutathiolation assessed using bioGEE (Fig 6C) and DTT-PEG exchange (Fig 6D) assays, although the glutathiolation status of C19A caveolin-3 was consistently reduced compared to wild type caveolin-3 in both assays. However, reintroduction of Cys-19 into cys-less caveolin-3 significantly increased caveolin-3 glutathiolation to a level not distinguishable from wild type (Fig 6E). We conclude that Cys-19 is the principal site of glutathiolation in caveolin-3, although we cannot rule out residual glutathiolation of other sites when this site is removed.

**Figure 6:**
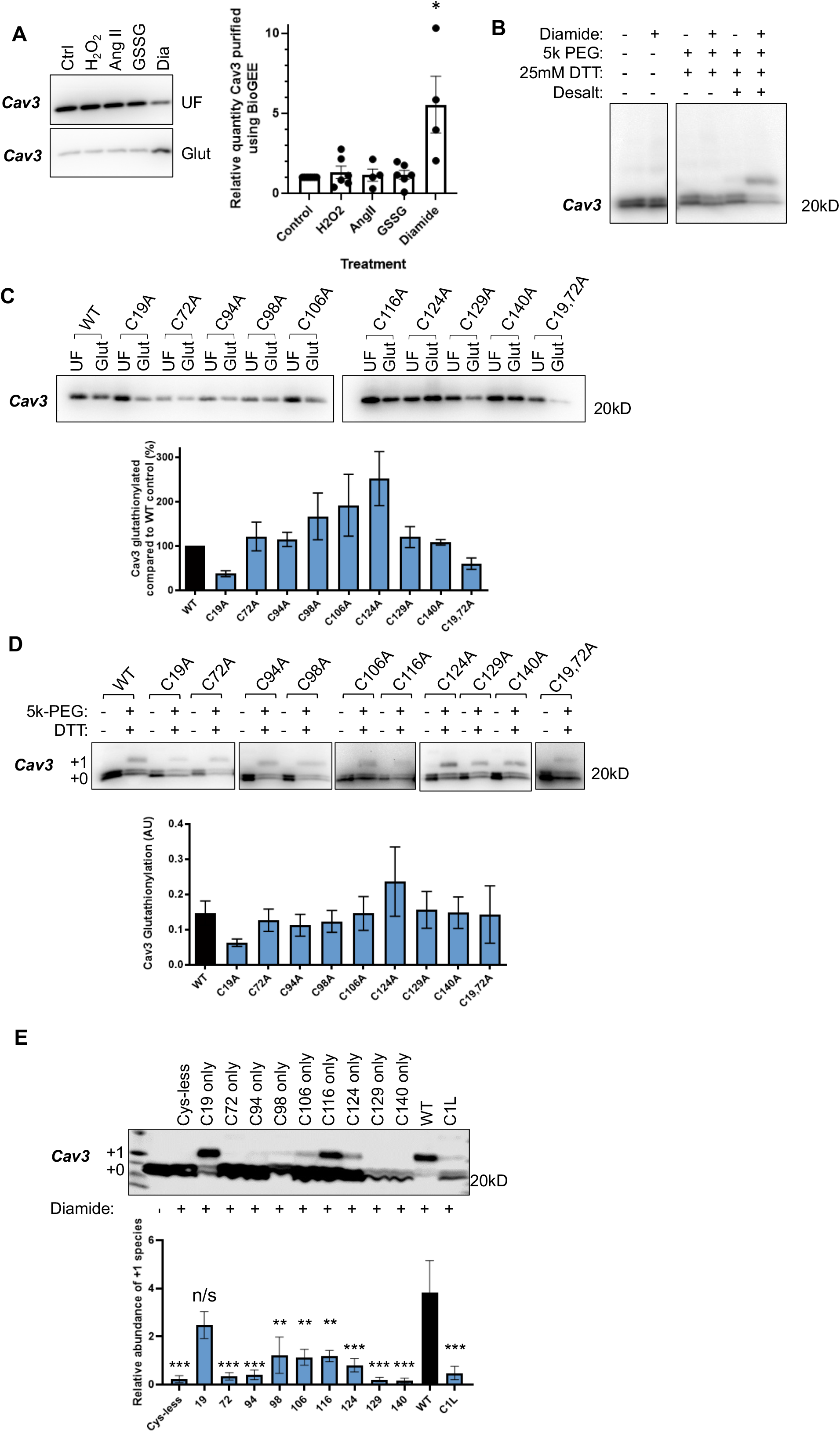
Glutathiolation of caveolin-3. A: Diamide treatment of rat ventricular myocytes increases glutathiolation of caveolin-3 assessed using bioGEE. B: PEGylation to measure caveolin-3 glutathiolation. Replacement of hydroxylamine with DTT in the acyl PEG exchange protocol reveals a diamide-induced bandshift of caveolin-3 consistent with a single glutathiolation site. C: Single cysteine to alanine mutagenesis of caveolin-3 does not significantly reduce caveolin-3 glutathiolation assessed using bioGEE (n=3-5, mean ± SEM shown). D: Single cysteine to alanine mutagenesis of caveolin-3 does not significantly reduce caveolin-3 glutathiolation assessed using DTT-dependent PEGylation (n=3-5, mean ± SEM shown). E: Only the reintroduction of Cys 19 into cys-less caveolin-3 restores glutathiolation to wild type levels. Abundance of the +1 glutathiolated band is expressed relative to the abundance of the +0 non-glutathiolated band. (C1L: caveolin-1 like; n=5; statistical comparisons shown to WT; **: P<0.01; ***: P<0.001; one way ANOVA followed by Dunnett’s multiple comparisons test).

### Functional Impact of Caveolin-3 Glutathiolation

Using diamide as a tool compound, we investigated the impact of increased caveolin-3 glutathiolation on caveolae in isolated ventricular myocytes. Oligomerisation of caveolin-3 can be visualised by partial formaldehyde fixation of intact cells, followed by electrophoresis and western blotting to detect oligomerised caveolins (22). We found no difference in caveolin-3 oligomerisation following diamide treatment (Fig 7A), nor did diamide change the gross distribution of caveolin-3 on sucrose gradients (Fig 7B). Indeed, when we considered the localisation of caveolin in buoyant (fractions 4 & 5) and dense (fraction 8) membranes isolated on a sucrose gradient, we detected no difference in the extent of palmitoylation or glutathiolation between fractions (Fig 7C). We conclude that glutathiolation of caveolin-3 Cys-19 does not substantially remodel caveolae, so instead we investigated caveolin-3 protein interactions.

**Figure 7:**
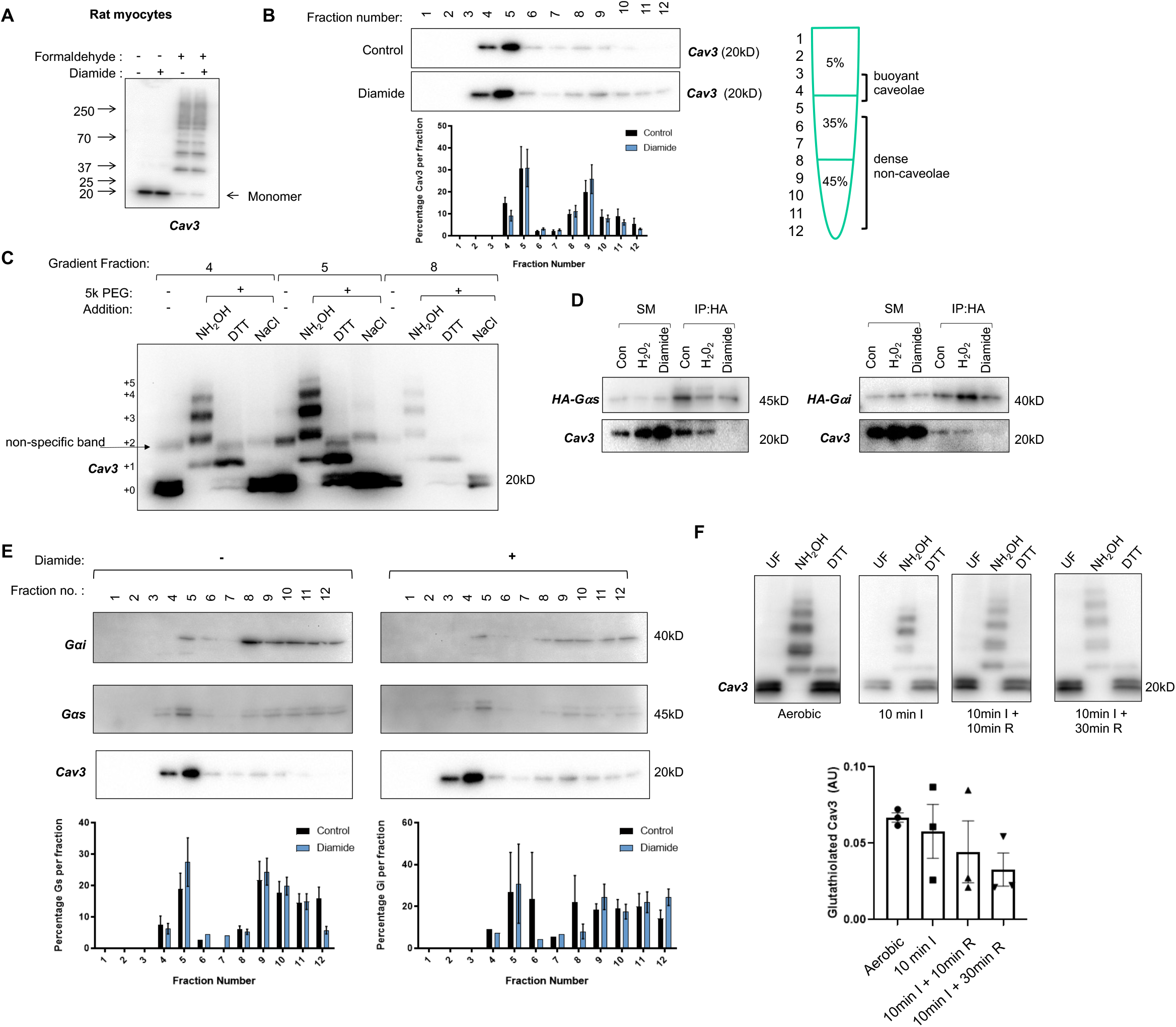
Impact of glutathiolation of caveolin-3. A: Partial formaldehyde fixation reveals caveolin-3 oligomerisation is not altered by diamide-induced glutathiolation of caveolin-3. B: Sucrose gradient fractionation reveals caveolin-3 distribution across sucrose gradient fractions is not altered by diamide-induced glutathiolation (n=5, mean ± SEM shown). C: PEGylation to investigate differential post-translational modifications of caveolin-3 in sucrose gradient fractions. Caveolin-3 is equally palmitoylated (NH_2_OH: hydroxylamine-dependent PEGylation) and glutathiolated (DTT: DTT-dependent PEGylation, NaCl: negative control) in buoyant and non-buoyant membranes. D: Co-immunoprecipitation of caveolin-3 and G protein subunits is abolished when caveolin-3 is glutathiolated. SM: starting material; IP: immunoprecipitation. E: Localisation of G protein subunits to buoyant, caveolin-enriched microdomains is not altered when caveolin-3 is glutathiolated (n=5, mean ± SEM shown). F: Neither caveolin-3 palmitoylation (NH_2_OH: hydroxylamine-dependent PEGylation) nor glutathiolation (DTT: DTT-dependent PEGylation) are changed during cardiac ischemia or following brief (10min) or long (30min) periods of reperfusion. 10min I: 10min global ischemia. 10min I + 10min R: 10min global ischemia followed by 10min reperfusion. 10min I + 30min R 10min global ischemia followed by 30min reperfusion.

Some proteins are recruited to caveolae as a result of their interaction with a juxtamembrane scaffolding domain in caveolin (CSD, Fig 1) (4). Although the concept of the CSD as a universal means to target proteins into caveolae has been challenged in recent years (12,23,24), interactions between the CSD and certain caveolar proteins (e.g. G-protein alpha subunits, tyrosine kinases) are well established, and generally inhibit the activity of these signalling proteins (25,26). We evaluated interaction of caveolin-3 with HA-tagged G protein alpha subunits in transfected HEK cells, and the impact of caveolin-3 glutathiolation on these interactions by treating with diamide (Fig 7D). Exogenous hydrogen peroxide, which does not alter caveolin-3 glutathiolation, did not change the co-immunoprecipitation of caveolin-3 and either Gαi or Gαs, but treatment with diamide abolished the binding of caveolin-3 to both G proteins. We conclude that glutathiolation of caveolin-3 Cys-19 reduces the ability of the caveolin-3 CSD to interact with its binding partners.

Given the impact of caveolin-3 gluathiolation on its protein interactions, we evaluated the impact of caveolin-3 glutathiolation on G protein alpha subunit localisation to caveolae using sucrose gradient fractionation of isolated adult rat ventricular myocytes. Lipidation of G protein alpha subunit N termini (S-palmitoylation and myristolyation of Gαi, N-palmitoylation and S-palmitoylation of Gαs) recruit these proteins to caveolae / lipid rafts (27,28). We observed no difference in the distribution of either G protein across a standard discontinuous sucrose gradient following glutathiolation of caveolin-3, suggesting these G proteins remain localised in caveolae following their dissociation from caveolin-3 (Fig 7E).

The burst of oxygen free radicals produced by mitochondria upon reperfusion of ischaemic cardiac tissue causes oxidative modifications of cardiac lipids and proteins (29,30). We assessed whether global ischemia with or without reperfusion changed the glutathiolation status of caveolin-3 in rat hearts perfused in the Langendorff mode. Since it is impossible to load whole hearts with bioGEE, these experiments employed the DTT-PEG exchange assay to monitor caveolin-3 glutathiolation. We investigated a short period of ischemia (10min) that stuns but does not irreversibly injure the myocardium (31), and reperfused for short (10min) and long (30min) periods. Caveolin-3 glutathiolation was not significantly altered following ischaemia-reperfusion (Fig 7F).

## Discussion

In this investigation we set out to characterise cysteine post-translational modifications of caveolin-3. We find that up to six cysteines in caveolin-3 are post-translationally palmitoylated and a single cysteine is reversibly glutathiolated. Glutathiolation of caveolin-3 directly influences its binding to G proteins.

In cardiac muscle 100% of caveolin-3 is palmitoylated, evidenced by its quantitative capture using assays such as acyl-RAC and acyl biotin exchange (32,33). The acyl-PEG exchange assay used in this investigation quantitatively PEGylates caveolin-3 in the sense that no ‘+0’ species is identified following the PEGylation reaction. However, quantitative PEGylation of every previously palmitoylated cysteine requires there to be no steric impairment of multiple PEGylation events, which we do not rule out - particularly when considering the attachment of 6x 5kDa PEG molecules within a short (46 amino acid) region of a 17.5kDa protein. We therefore suggest that the predominant +4 palmitoylation stoichiometry for caveolin-3 from ventricular muscle identified by acyl-PEG exchange must be regarded as the ‘minimum’ rather than ‘absolute’ stoichiometry of caveolin-3 palmitoylation.

The cryo-EM structure of the caveolin-1 oligomer suggests that rather than possessing a classic re-entrant integral membrane loop, the oligomer displaces the cytosolic leaflet of the membrane bilayer, making direct contact with the outer leaflet of the membrane, into which the caveolin palmitates presumably insert (7). It is not currently understood how the profoundly flat caveolin oligomer can bend the membrane to induce the characteristic caveolar shape. This shape is acquired even when caveolin-1 is expressed in bacteria (19), where it is not palmitoylated, so palmitoylation is clearly not integral to this process. Nevertheless, the ability of heavily palmitoylated proteins to induce membrane curvature and substantially remodel their local phospholipid environment is well established (34,35). For example, molecular dynamics simulations suggest palmitoylation of the SARS-CoV-2 spike protein (30 palmitates per spike trimer) induces formation of ordered cholesterol and sphingolipid-rich microdomains (36). As well as modifying its phospholipid environment, we predict that the presence of up to 66 palmitates per caveolin-3 oligomer will significantly alter the packing of the phospholipid tails in the outer leaflet of the bilayer compared to the influence of up to 33 palmitates for oligomeric caveolin-1. The unique phospholipid environment in caveolae is achieved by lipid interactions of the caveolar structural proteins (37,38). Our results therefore provide a mechanistic basis by which the phospholipid composition and overall architecture of caveolin-3 containing caveolae could be distinct from those formed from caveolins 1 or 2.

What are the consequences of differential caveolin isoform palmitoylation for caveolar resident proteins? Although caveolin-3 is regarded as the muscle-specific caveolin isoform, all three caveolin isoforms are expressed in ventricular muscle, where caveolins 1 and 3 localise to distinct but adjacent microdomains (39). The protein interactions of caveolin-1 and caveolin-3 in cardiac muscle are different. For example, caveolin-1 preferentially interacts with cavins 1 and 2 and aquaporin-1, whereas caveolin-3 interacts with transporter proteins such as NCX1, Na/K ATPase, Glut-4 and the monocarboxylate transporter McT1 (12,39). If the differential palmitoylation profile of caveolin isoforms drives the formation of distinct phospholipid microenvironments, different protein species are likely to be attracted to these microenvironments. Ultimately then, proteins may be either colocalised or kept apart in separate microdomains as a result of the differential lipid post-translational modifications of the caveolins that form these domains.

Considering the positions of the additional palmitoylation sites in the caveolin-3 oligomer, two additional concentric ‘rings’ of palmitoylated cysteines exist compared to caveolin-1. We did not detect significant palmitoylation of the cysteine closest to the 11-stranded β-barrel located at the centre of the structure, but in contrast the outermost ring of palmitoylated cysteines (Cys-94, Cys-98) adds palmitate at the edge of the oligomer, where phospholipid interactions may occur. In this context it is notable that Cys-72 from one protomer is immediately adjacent to Cys-94 in the adjacent protomer but remains non-palmitoylated. The caveolin oligomer assembles in the endoplasmic reticulum (40) but is palmitoylated later in the secretory pathway (16). Evidently there is sufficient specificity to caveolin-3 palmitoylation to prevent palmitoylation at Cys-72 despite its close proximity to other palmitoylated cysteines. How the active site of the caveolin-3 palmitoylating enzyme(s) manages to access up to 66 cysteines across the hydrophobic face of the assembled caveolin oligomer requires further investigation.

The caveolin scaffolding domain proposed to interact with caveolar resident proteins is residues 61-101 of caveolin-1 (34-74 of caveolin-3) (25). Caveolin-1 82-101 directly interacts with heterotrimeric G Protein α subunits (4), but we found caveolin-3 interaction of G protein α subunits was reduced when it was glutathiolated at Cys-19, which lies outside the proposed caveolin scaffolding domain. This region of the caveolin N terminus is disordered and ripe for protein-protein interactions. Post-translational modifications, even outside the site of interaction, may cause the caveolin N terminus to adopt a conformation that does not interact favourably with G proteins. No matter how Cys-19 glutathiolation abolishes caveolin-3’s interaction with G proteins, it is notable that this redox sensitivity of protein interaction is another unique adaptation of the muscle-specific caveolin isoform identified in this investigation. Since the interaction of caveolins with G proteins is generally regarded as inhibitory, our results suggest a paradigm in which redox stress can modify signalling downstream of G proteins. A similar paradigm is already established for the kinases regulated by second messengers produced by these G proteins (41,42). An important question for future investigations will be how glutathiolation of caveolin-3 modifies G protein activation and second messenger production.

## Methods

### Ethics statement

All procedures were approved by University of Dundee Animal Welfare and Ethical Review Board. The animal research reported here adheres to the ARRIVE and Guide for the Care and Use of Laboratory Animals guidelines.

### Plasmids and Cells

Rat caveolin-3 in pExpress-1 (IMAGE ID 7103841) was purchased from Source Bioscience. Site mutants of caveolin-3 were generated using the Agilent Quikchange II and Quikchange Lightning Multi site directed mutagenesis kits using oligonucleotide primers designed according to the manufacturer’s recommendations.

HEK-293 and H9c2 cells were cultured in DMEM supplemented with 10% fetal calf serum. HEK-293 cells were transfected using Lipofectamine 2000 in 6-well and 12-well culture plates and harvested 18-24 hours after transfection.

Calcium-tolerant rat ventricular myocytes were isolated by retrograde perfusion of collagenase in the Langendorff mode, as described previously (43).

Cultured cells and ventricular myocytes were treated with hydrogen peroxide (100μM, 10-30min), angiotensin II (500nM, 20min), oxidised glutathione (GSSG 5mM, 10min) or diamide (1mM, 10-15min) before lysis.

### Langendorff Heart

Hearts from adult male Wistar rats (250-300g, Charles River Laboratories) were perfused in the Langendorff mode at a constant pressure of 100cm water using Krebs-Henseleit buffer equilibrated with 95% oxygen, 5% CO2 at 37°C. Global ischemia was induced by complete cessation of coronary flow, with normothermia during ischemia ensured by immersing the heart in Krebs-Henseleit during the ischemic period.

### SDS PAGE and Western Blotting

Proteins were separated on 6-20% gradient gels, transferred to PVDF membranes and incubated with primary antibodies overnight at 4°C. Western blot images were acquired using a ChemiDoc XRS imaging system and quantified using QuantityOne (BioRad), or using a LICOR Odyssey Fc and quantified using ImageStudio (LICOR). When quantifying PEGylation, we measured the abundance of the PEGylated species relative to the abundance of the non-PEGylated species of the protein in the same lane.

### Palmitoylation assays

We adapted our PEG-switch assay that replaces palmitate with a 5kDa methoxypolyethylene glycol (PEG) (44) using a refinement that improves the reaction efficiency by PEGylating separately from the thioester cleavage step (45,46). Briefly, free protein thiols were alkylated with 100mM maleimide (in the presence of 2.5% SDS, 100mM HEPES, 1mM EDTA, pH 7.5) for 4hours at 40°C, and excess maleimide removed by acetone precipitating proteins and extensive washing of the protein pellet with 70% acetone. Proteins were resolubilised (1% SDS, 100mM HEPES, 1mM EDTA, pH7.5) and thioester bonds cleaved by treating with 200mM neutral hydroxylamine for 1hour at 37°C. Hydroxylamine was removed by desalting (Zeba spin column) and free cysteines PEGylated with 2mM 5K-PEG maleimide (Sigma) for 1hour at 37°C.

Palmitoylated proteins were purified from whole cell lysates using acyl-resin assisted capture (Acyl-RAC). Briefly, free thiols were alkylated with methyl methanethiosulfonate and palmitoylated proteins captured using thiopropyl-Sepharose in the presence of neutral hydroxylamine (47).

### Glutathiolation assays

We synthesised biotinylated glutathione ethyl ester (bioGEE) by reacting sulfo-NHS-biotin (Thermo) with glutathione ethyl ester (Sigma) in 50mM NaHCO3 pH 8.5 for 2 hours at room temperature. Unreacted sulfo-NHS-biotin was quenched with 125mM NH4HCO3 for 1 hour at room temperature. Cells were loaded by incubating with 500μM bioGEE for 1 hour then washed twice prior to experimentation. Cells were lysed with 1% Triton X-100 in PBS supplemented with protease inhibitors, biotinylated proteins captured with streptavidin-Sepharose for 1hour at 4°C and eluted using 100mM DTT after extensive washing.

For mass spectrometry analysis, samples were alkylated with iodoacetamide, concentrated using 10kDa cut-off centrifugal filters, digested with trypsin overnight, acidified, and cleaned up using C18 zip-tips. 0.3-0.5μg of each sample was analysed using a nanoflow liquid chromatograph (Agilent 1200, Agilent, Santa Clara, CA) with an LTQ-Orbitrap XL (Thermo Fisher Scientific). Protein and peptide database search was carried out using PEAKS version 6 (Bioinformatics Solutions, Waterloo, Canada). A minimum of two unique peptides were required for each protein identified.

We adapted our PEG-switch assay to assess protein glutathiolation. Reactions proceeded exactly as for acyl-PEG exchange, with the exception that hydroxylamine was replaced with 25mM DTT, which was applied at 37°C for 10min before desalting and treating with 2mM 5K-PEG maleimide for 1hour at 37°C.

### Sucrose Gradient Fractionation

We used a standard discontinuous sucrose gradient to separate cholesterol-rich buoyant membranes from bulk sarcolemma and intracellular membranes following homogenisation and sonication of whole cell lysates in 500mM sodium carbonate (12).

### Partial Formaldehyde Fixation

Caveolin-3 oligomers were visualised by partial fixation in 1% formaldehyde in PBS for 20min followed by electrophoresis and immunoblotting (22).

### Co-immunoprecipitation

HA-tagged and EE-tagged G protein alpha subunits were immunoprecipitated from HEK cell lysates as described previously (44).

## Acknowledgements

We acknowledge the support of the British Heart Foundation. FS/13/22/30126 to WF and CH, PG/15/42/31563 to WF, SC and IJ, PG/19/5/34150 to WF.

